# Performance of gene expression analyses using *de novo* assembled transcripts in polyploid species

**DOI:** 10.1101/380063

**Authors:** Ling-Yun Chen, Diego F. Morales-Briones, Courtney N. Passow, Ya Yang

## Abstract

**Motivation:** Quality of gene expression analyses using *de novo* assembled transcripts in species experienced recent polyploidization is yet unexplored.

**Results:** Five plant species with various polyploidy history were used for differential gene expression (DGE) analyses. DGE analyses using putative genes inferred by Trinity performed similar to or better than Corset and Grouper in precision, but lower in sensitivity. In species that lack polyploidy event in the past few million years, DGE analyses using *de novo* assembled transcriptome identified 50–76% of the differentially expressed genes recovered by mapping reads to the reference genes. However, in species with more recent polyploidy event, the percentage decreased to 7–30%. In addition, 7–89% of differentially expressed genes from *de novo* assembly are contaminations. Gene co-expression network analyses using *de novo* assemblies vs. mapping to the reference genes recovered the same module that significantly correlated with treatment in one of the five species tested.

**Availability and Implementation:** Commands and scripts used in this study are available at https://bitbucket.org/lychen83/chen_et_al_2018_benchmark_dge/; Analysis files are available at Dryad doi: XXXXXX.

**Supplementary information:** Supplementary data are available at *Bioinformatics* online

## 1 Background

With decreasing sequencing cost, *de novo* assembled transcriptomes have been increasingly used for exploring gene space and gene expression in diverse species without reference genome (Misof *et al*., 2014; Heyduk *et al*. 2018). This approach has been especially valuable in plants, where genome sizes are relatively large compared to fungi or animals. Because of active development of sequencing platforms and *de novo* assembly tools, *de novo* assembled transcriptomes can recover up to 75% of genes in plant genomes (Honaas *et al*., 2016) and 78% in animals (Carruthers *et al*., 2018). The improved coverage and accuracy make downstream analyses, such as differential gene expression, phylotranscriptomics, and gene co-expression networks possible based on *de novo* transcriptome assemblies.

Despite the improvements, key issues still remain. These include clustering assembled transcripts into putative genes, removing redundant transcripts that are often assembly artifacts, and obtaining a representative transcript for each putative gene. These steps are important because differential gene expression (DGE) analysis is more accurate and interpretable than differential transcript expression (DTE) analysis (Soneson *et al*., 2015; Davidson and Oshlack, 2014). Several strategies have been proposed to cluster assembled transcripts to putative genes. *De novo* transcriptome assemblers such as Trinity (Grabherr *et al*., 2011) and Oases (Schulz *et al*., 2012) cluster transcripts into putative genes based on de Bruijn graph structure during assembly. On the other hand, assembler-independent clustering approaches, such as CD-HIT-EST (Li and Godzik, 2006), Corset (Davidson and Oshlack, 2014) and Grouper (Malik *et al*., 2018) allow transcripts from any source to be clustered after the assembly stage. Previous benchmark studies comparing transcript clustering methods yielded contradictory results. For example, the performances of CD-HIT-EST and Corset in Davidson and Oshlack (2014) and Srivastava *et al*. (2016).

In addition to the conflicting results, most benchmark studies of DGE analysis using *de novo* assembled transcriptome used data from animals and fungi with only a few using plants (e.g. Wang and Gribskov, 2017; Malik *et al*., 2018). In addition, previous studies have avoided species with recent polyploidy history. Recent comparative genomic and transcriptomic analyses have revealed occurrence of many ancient and more recent whole genome duplication events in insects (Li *et al*. 2018), vertebrates (Berthelot *et al*. 2014), fungi (Marcet-Houben and Gabaldón, 2015), and much more frequently in plants (Lee *et al*., 2017). The timing of the last round of polyploidization can impact *de novo* transcriptome assembly and any downstream analysis (Nakasugi *et al*., 2014). However, to what extent the impact remains to be quantified.

With the further decrease in costs for transcriptome sequencing, gene co-expression network analysis has been gaining popularity in the past few years. The analysis was used to identify genes associated with traits such as disease, metabolites, and stress response (Loraine, 2009; Garcia *et al*., 2017). With a few exceptions (e.g. Heyduk *et al*., 2018; Roberts and Roalson, 2017), the analysis was limited to species with the reference genomes available. The reliability of carrying out such co-expression analysis using *de novo* transcriptome assemblies has never been evaluated, limiting our ability to dissect the genetic basis of complex traits in diverse organisms without reference genomes.

Using two species with ancient polyploidy events only, and three with more recent polyploidy events, here we evaluate three important issues in downstream analyses of *de novo* assembled transcripts: 1) performance of methods for clustering transcripts into putative genes; 2) performance of DGE analyses in polyploid species; and 3) whether gene co-expression networks recovered from *de novo* assemblies are comparable with those recovered from reference-based analysis.

## 2 Materials and Methods

### 2.1 Data and *de novo* transcriptome assembly

In this study we chose five plant species that 1) each has a well-annotated reference genome; 2) each has at least 12 publicly available RNA-seq datasets; 3) each data point has at least three biological replicates for differential expression analysis; and 4) vary in age of the last polyploidy event (Table 1). In total, five species were selected. Both Arabidopsis (*Arabidopsis thaliana*) and grape (*Vitis vinifera*) are ancient polyploids, with the most recent polyploidy events being 65.5 and 118 million years ago respectively (Ma; Beilstein *et al*., 2010; Forest and Chase, 2009; Lee *et al*., 2012). Maize (Zea mays), on the other hand, experienced a more recent polyploidy event at 4.8-15.4 Ma (Blanc and Wolfe, 2004; Swigoftová *et al*., 2004). Rapeseed (*Brassica napus*) is an allotetraploid with the split between the two parental genomes at approximately 2.5 Ma (Arias *et al*., 2014). The common wheat (*Triticum aestivum*) is an allohexaploid with the split between parental genomes at approximately 2 Ma (Middleton *et al*., 2014). See Supplementary Table S1 for additional details

Non-redundant transcripts (referred to “primary transcripts” in Phytozome), which do not include splice variants, were used as “reference genes” for benchmark analyses. To verify the polyploidy history for each species, we visualized the distribution of synonymous substitutions (*Ks*) for within-species paralog pairs using reference genes with the script ks_plots.py (Yang *et al*., 2015). Default parameters were used for all analyses throughout this study unless noted otherwise. Random sequencing errors were corrected using Rcorrector v.1.0.2 (Song and Florea, 2015). Sequencing adaptors and low-quality bases were trimmed with Trimmomatic v.0.36 (Bolger *et al*., 2014) (TruSeq_adapters:2:30:10 SLIDINGWINDOW:4:15 LEADING:5 TRAILING:4 MINLEN:80). *De novo* transcriptome assembly for grape was carried out using Trinity v.2.6.5 (Grabherr *et al*., 2011). The remaining species were assembled by Trinity v.2.6.6 with minor updates including python-3 compatibility and update to R command execution.

**Table 1.**
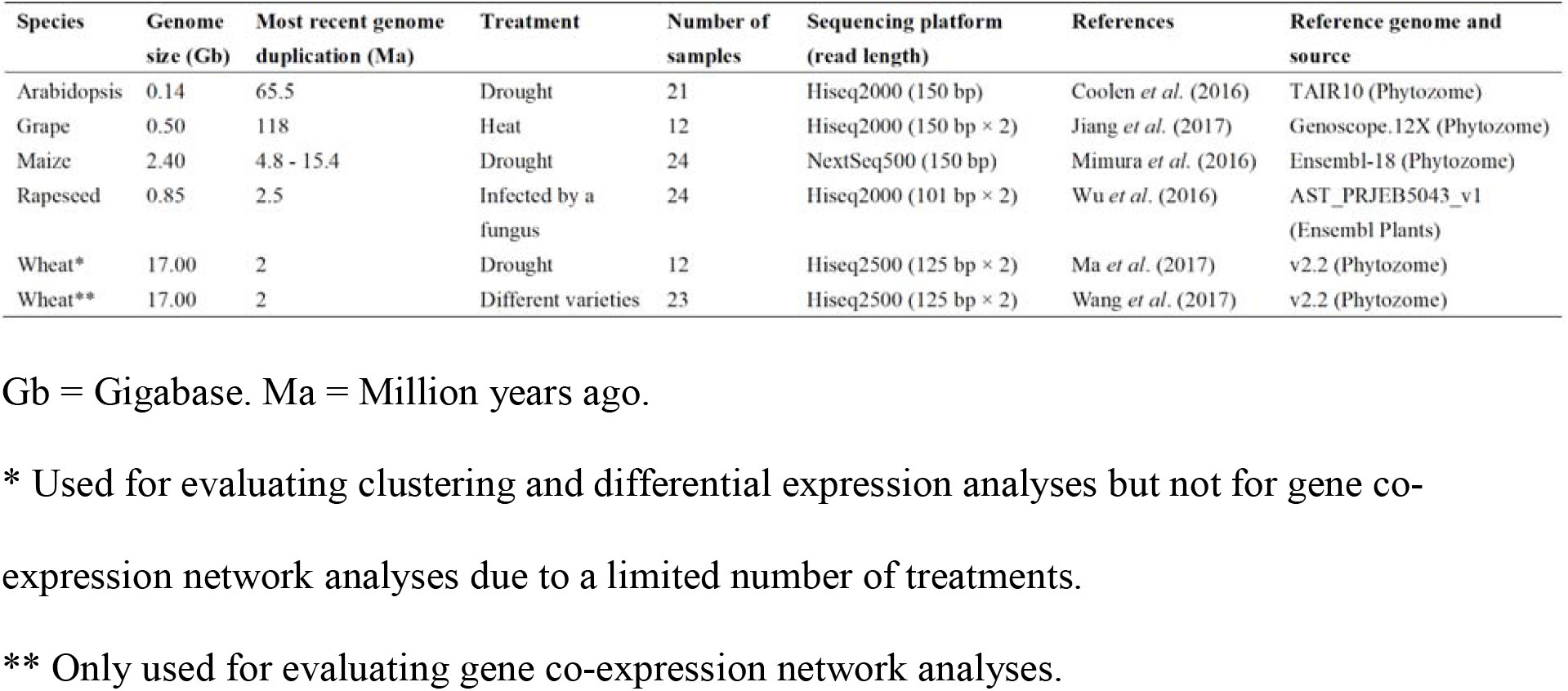
Summary of species, RNA-seq datasets, and reference genomes used in this study.

Given that decontamination and filtering non-coding sequences did not improve DGE analyses (Supplementary Methods and Supplementary Table S2 and Fig. S1), these procedures were not carried out in remaining species.

### 2.2 Clustering transcripts into putative genes

In this study, “clustering” refers to grouping *de novo* assembled transcripts that are likely to belong to the same gene (Malik *et al*., 2018). We used four strategies for clustering, Trinity, CD-HIT-EST, Corset, and Grouper. Trinity cluster information was directly extracted from sequence identifiers of assembled transcripts. CD-HIT-EST clusters transcripts by overall similarity. Assembled transcripts were clustered using CD-HIT-EST v.4.7 (Li and Godzik, 2006). For Corset (Davidson and Oshlack, 2014), reads from each sample were first mapped to assembled transcripts using “pseudo-alignment” method Salmon v.0.91 (Patro *et al*., 2017). Fragment equivalence classes generated by Salmon were used as input for Corset v.1.07. Corset groups all transcripts that share one or more reads into a super-cluster, and then use a hierarchical clustering algorithm to separate each super-cluster to clusters according to expression level. By doing so Corset is capable of teasing apart transcripts from different genes and chimeric transcripts (Davidson and Oshlack, 2014). Default parameters of Corset were first used. To prevent transcripts within a gene with differential transcript usage from splitting into different clusters, the parameter ‘-D 99999999999’ was used. By default, Corset ignores transcripts with less than ten reads mapped. We applied the parameter ‘-m 0’ to keep all transcripts. In order to explore the potential effect of different alignment tools, alignment method Bowtie v.2.3.4.1 (Langmead and Salzberg, 2012) was also used Corset in Arabidopsis. Since Salmon produced comparable mapping results to Bowtie2 and is more time and memory efficient (Teng *et al*., 2016), we carried out the read mapping only with Salmon for the remaining four species. Finally, the fourth strategy uses fragment equivalence classes from Salmon as input for clustering with Grouper v.0.1.1 (Malik *et al*., 2018). Grouper is similar to Corset except that the former uses the Markov cluster algorithm while the latter uses a hierarchical clustering algorithm.

### 2.3 Evaluating clustering methods

To determine the correspondence between assembled transcripts and reference genes, transcripts were aligned to reference genes using BLAT v.35 (Kent, 2002). Hits with match length ≥200 bp and identity ≥ 98% were considered as positive matches. In cases when a transcript had multiple positive matches, the gene with the longest match was kept.

We evaluated three aspects of the performance of clustering methods. First, we estimated total gene coverage. The longest transcript for each cluster was chosen as the representative transcript. Total gene coverage (number of reference genes with at least one positive match/total number of reference genes) was calculated after removing chimeric transcripts following Yang and Smith (2013).

Next, we evaluated precision, recall, and correct clusters of clustering *de novo* transcripts. Any two transcripts in the same cluster that each had a positive match to reference genes were evaluated for whether they were correctly placed in the same cluster (true positive, TP), correctly separated in different clusters (true negative, TN), incorrectly placed in the same cluster (false positive, FP), or incorrectly separated in different clusters (false negative, FN; Davidson and Oshlack, 2014). Precision = TPs / (TPs + FPs), recall = TPs / (TPs + FNs) (Davidson and Oshlack, 2014). F-score, which balances the precision and recall was calculated by 2 × precision × recall / (precision + recall) (van Rijsbergen, 1979). In addition, the number of correct clusters was calculated for each species. For a cluster to be “correct”, we required all *de novo* assembled transcripts in the cluster best matched to the same reference gene, and the cluster must include all transcripts that were best matched to the reference gene. Transcripts failed the above BLAT filter were ignored. Therefore, if none of the assembled transcripts within a cluster had a positive match to any reference gene, the cluster was ignored for evaluating performance of clustering.

Last, the impact of clustering approaches in differential expression analyses was evaluated by either picking the longest transcript within a cluster to represent a putative gene for read mapping, or summarizing all reads mapped to all transcripts in a cluster. Each cluster was matched to its corresponding reference gene according to the best match by the majority of its transcripts with the same filter as above. When an equal number of transcripts within a cluster matched to multiple reference genes, the gene with the longest aligned length was selected. In addition to DGE analyses at the gene level, DTE analyses were carried out by mapping reads to all transcripts from *de novo* assembly without taking clusters into consideration. For both strategies, Salmon was used to quantify expression level. In addition, RSEM v.1.3 (Li and Dewey, 2011) with STAR v.2.5.3 (Dobin *et al*., 2013) were used to quantify expression level in grape to explore potential biases from mapping software packages.

After read mapping, differential expression analyses were conducted using DESeq2 v.1.20.0 (Love *et al*., 2014). Tximport v.1.6.0 (Soneson *et al*., 2015) was used to import expression data from Salmon or RSEM to DESeq2. Given that filtering putative genes with low expression level only slightly increased precision and recall, but decreased sensitivity (Supplementary Methods and results and Supplementary Fig. S1), outputs derived from DESeq2 without the filtering were used to evaluate clustering for all five species. |log2FoldChange| ≥ 2 and p-value ≤ 0.05 were used as the criterion of differential expression. DTE analyses using all assembled transcripts were carried out using the same settings.

Differentially expressed genes using *de novo* assembly (deDEG) were compared against analyses based on mapping to the reference genes. If a deDEG matched to a DEG from reference-based analysis (refDEG), it was considered as a TP; if matched to a reference gene that was not differentially expressed, as a FP. If a transcript was not a deDEG but a refDEG, it was considered as a FN; otherwise, if a transcript was neither a reDEG nor a deDEG, it was considered as a TN. We only included assembled transcripts that had a positive match to the reference genes for counting the number of TPs, TNs, FPs, and FNs. DESeq2 results were sorted by p-values, and the transcript with the lowest p-value was ‘unique’ when more than one transcript was matched to a reference gene. Precision, recall and F-score for differential expression analyses were calculated using unique true positives (UTPs), unique false positives (UFPs) and unique false negatives (UFNs) using the same formulas as for evaluating clustering. Sensitivity = UTPs / total positives from reference. Unique true positive rate (UTPR) = UTPs / total unique positives from *de novo* analysis, unique false positive rate (UFPR) = unique false positives (UFPs) / total unique negatives from *de novo* analysis (Fawcett, 2006). Receiver operating characteristic (ROC) curve was plotted by using corresponding UFPR and UTPR with 34 p-values from 0.0 to 1.0. Area under curve (AUC) was calculated by Σ(UFPR_n_ – UFPR_n−1_) × (UTPR_n_ + UTPR_n−1_) / 2. When n = 1, UFPR_n−1_ and UTPR_n−1_ = 0. For evaluating DGE results recovered from summarizing reads to gene level and DTE results, same formulas were used.

To explore the composition of all the deDEGs, we aligned them against their corresponding reference genomes and annotations using rnaQUAST v.1.5.1 (Bushmanova *et al*., 2016) with BLAT (minimum sequence identity = 90%). The deDEGs did not have any match were searched against a local NCBI non-redundant protein sequences database (NR; downloaded February 1, 2018) using BLASTX in NCBI blast+ v.2.2.29 (Camacho *et al*., 2009).

### 2.4 Gene co-expression network analysis

Weighted gene co-expression network analysis was conducted using WGCNA v.1.63 (Langfelder and Horvath, 2008). Reads were mapped to the longest transcript of each Trinity cluster using Salmon (Patro *et al*., 2017). Transcripts per million (TPM) from Salmon were used as input for WGCNA. The function *goodSamplesGenes* in WGCNA was used to detect genes and samples with elevated numbers of missing values. Samples for each species were clustered using the function *hclust*. The resulting dendrograms were visualized and outlier samples were removed. Gene network for each species was constructed using the function *blockwiseModules*. Minimum module size was set to 30. Treatment information was used as trait data (Supplementary Table S1) except wheat, in which the spike complexity data were used as traits (Wang *et al*., 2017). Hub genes were identified with kME >0.9. Modules represented by hub genes were correlated with traits, and the P-value was calculated using the function *corPvalueStudent*. Correlation value ≥0.8 and p-value ≤0.05 were treated as significantly correlated. Gene co-expression network analyses were also conducted by mapping to reference genes with the same settings. Hub genes from modules recovered from *de novo* assemblies were compared to those from reference genes by BLAT searches. The reference module that contains the highest number of genes matched to a *de novo* module was identified.

## 3 Results and discussion

In this study, we tested four strategies of clustering assembled transcripts into putative gene, and quantified the performance of these strategies in downstream differential expression analyses. We found a consistent trend that when genomes become more complex, the performance of both clustering and differential expression analyses become worse, despite datasets from the five species were generated using different versions of Illumina HiSeq platforms and the treatments and sequencing depths are different. We also found that due to the tendency of Trinity to over clustering compared to both Corset and Grouper, clusters produced by Trinity have higher precision but lower sensitivity in downstream DGE analyses. CD-HIT-EST, on the other hand, performed the worst in all aspects. Co-expression network analysis using assembled transcripts recovered similar results compared to reference-based analysis in Arabidopsis, but not in any remaining species.

### 3.1 *De novo* transcriptome assembly recovered half to two-thirds of genes in the reference genome

*De novo* transcriptome assembly using Trinity recovered 63,557, 271,810, 230,920, 318,046 and 353,181 transcripts for Arabidopsis, grape, maize, rapeseed, and wheat respectively. Of them, 1.2-8.3% were chimeric. After removing chimeric transcripts, *de novo* assembly recovered 64.1%, 66.5%, 45.8%, 46.6% and 47.4% of genes in the reference genome (Supplementary Table S3). When using the longest transcript in each cluster as representative, reference gene coverage decreased by approximately 5–10% in Arabidopsis and grape (Supplementary Table S3). The reference coverage decreased even more, by 10–20% in maize, rapeseed, and wheat, all of which are recent polyploids.

*Ks* plots confirmed that both Arabidopsis and grape lack any recent polyploidy event, whereas maize, rapeseed, and wheat had recent polyploidy events, with the most recent *Ks* peaks at 0.2 or lower (Supplementary Fig. S2).

### 3.2 Grouper and Corset slightly outperformed Trinity in clustering transcripts into putative genes, and CD-HIT-EST performed the worst among the four

Among the four clustering strategies, CD-HIT-EST recovered the highest numbers of clusters. However, it recovered the fewest number and the lowest proportion of correct clusters (Fig. 1 and Supplementary Table S3). Corset and/or Grouper produced higher numbers of correct clusters than Trinity in all five species except grape. The proportion of correct clusters was the highest in Arabidopsis, accounting for 51–59% of the evaluated clusters when using Trinity, Corset or Grouper. The proportion decreased to 35–50% in grape and further decreased to approximately 22% in maize, rapeseed, and wheat (Fig. 1 and Supplementary Table S3).

**Figure 1.**
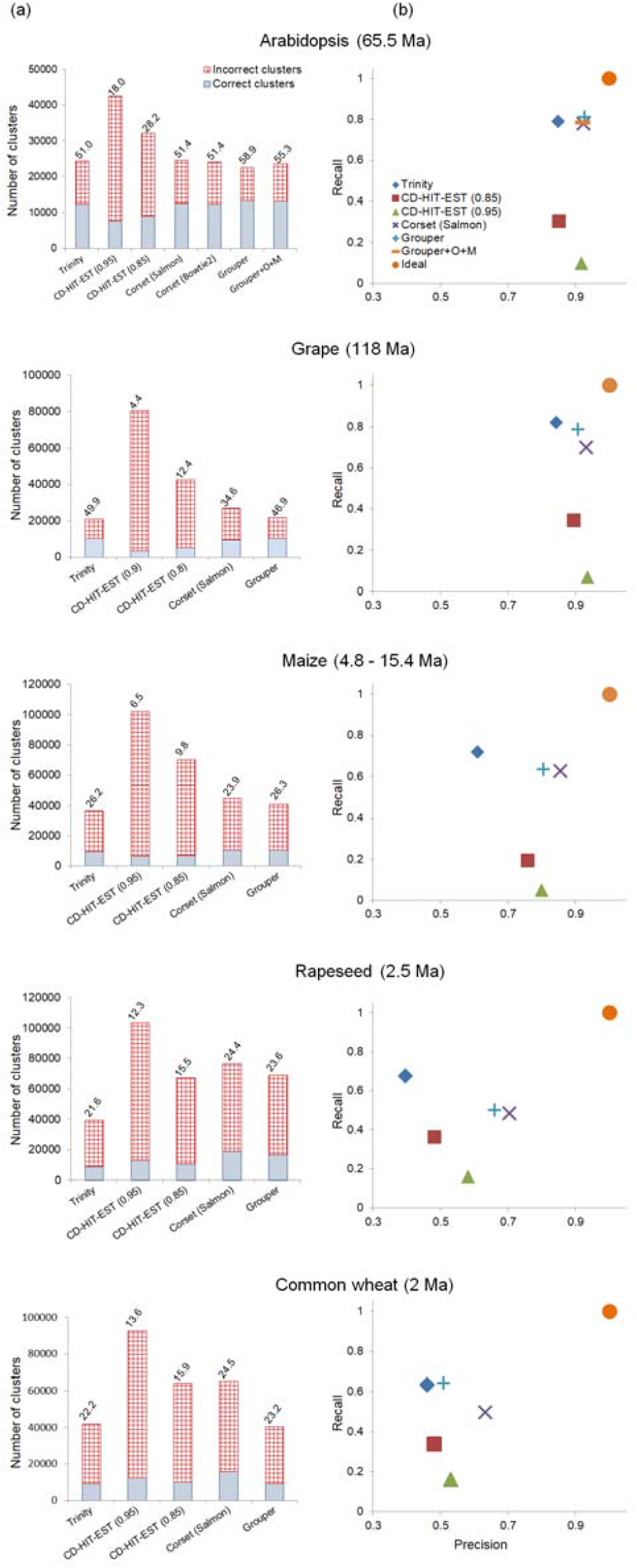
Performance of clustering methods. (a) Number and percentage (above the bars) of correct clusters. (b) Evaluating clustering by pairwise combinations of all transcripts in a given cluster. Sequence identity thresholds of 0.9 and 0.8 were used for the CD-HIT-EST analyses in grape only.

Among the four clustering methods, Trinity performed the best in recall, whereas CD-HIT-EST was the worst in recall (Fig. 1b). Grouper is similar or slightly higher in recall but lower in precision compared to Corset, consistent with Malik *et al*. (2018). Corset and Grouper had the best balance between precision and recall as shown by F-scores. Given that low recall indicates under clustering (splitting transcripts from the same gene to different clusters) and low precision indicates over clustering (grouping transcripts from different genes into the same cluster; Davidson and Oshlack, 2014), our study shows that Trinity tends to over clustering, while Corset, Grouper, and especially CD-HIT-EST tend to under clustering, and the difference is increasingly pronounced in more recent polyploids (Fig. 1).

Among the five species investigated, precision and recall were the highest in Arabidopsis with the F score ranging from 0.819 to 0.865 for Trinity, Corset and Grouper, followed by grape and maize (Supplementary Table S3). Precision and recall were low in rapeseed and wheat, with F scores ranging from 0.498 to 0.567.

### 3.3 Clusters generated by Trinity have higher precision but lower sensitivity than those from Corset and Grouper in downstream differential gene expression analyses

Compared to DGE analyses, DTE yielded lower precision, without increasing recall or sensitivity (Supplementary Table S4), which is consistent with Davidson and Oshlack (2014). The lower precision may come from a low number of UTPs and/or a high number of UFPs. In DTE analysis, reads may be ambiguously assigned to multiple transcripts in a gene, leading to incorrectly identifying differentially expressed transcripts (Soneson *et al*., 2015).

When representing a gene using the longest transcripts in DGE analysis, Trinity had higher precision but lower sensitivity than Corset and Grouper (Supplementary Table S4). Given that Trinity tends to over clustering compared to both Corset and Grouper, it produces a smaller number of UTPs but a much smaller number of UFPs. For example, in grape, while Corset recovered 149 UTPs and 156 UFPs, Trinity recovered 130 UTPs and 70 UFPs. ROC curve showed that the AUC value of Trinity was slightly higher than Corset and Grouper, and much higher than CD-HIT-EST (Fig. 2c and Supplementary Table S4).

**Figure 2.**
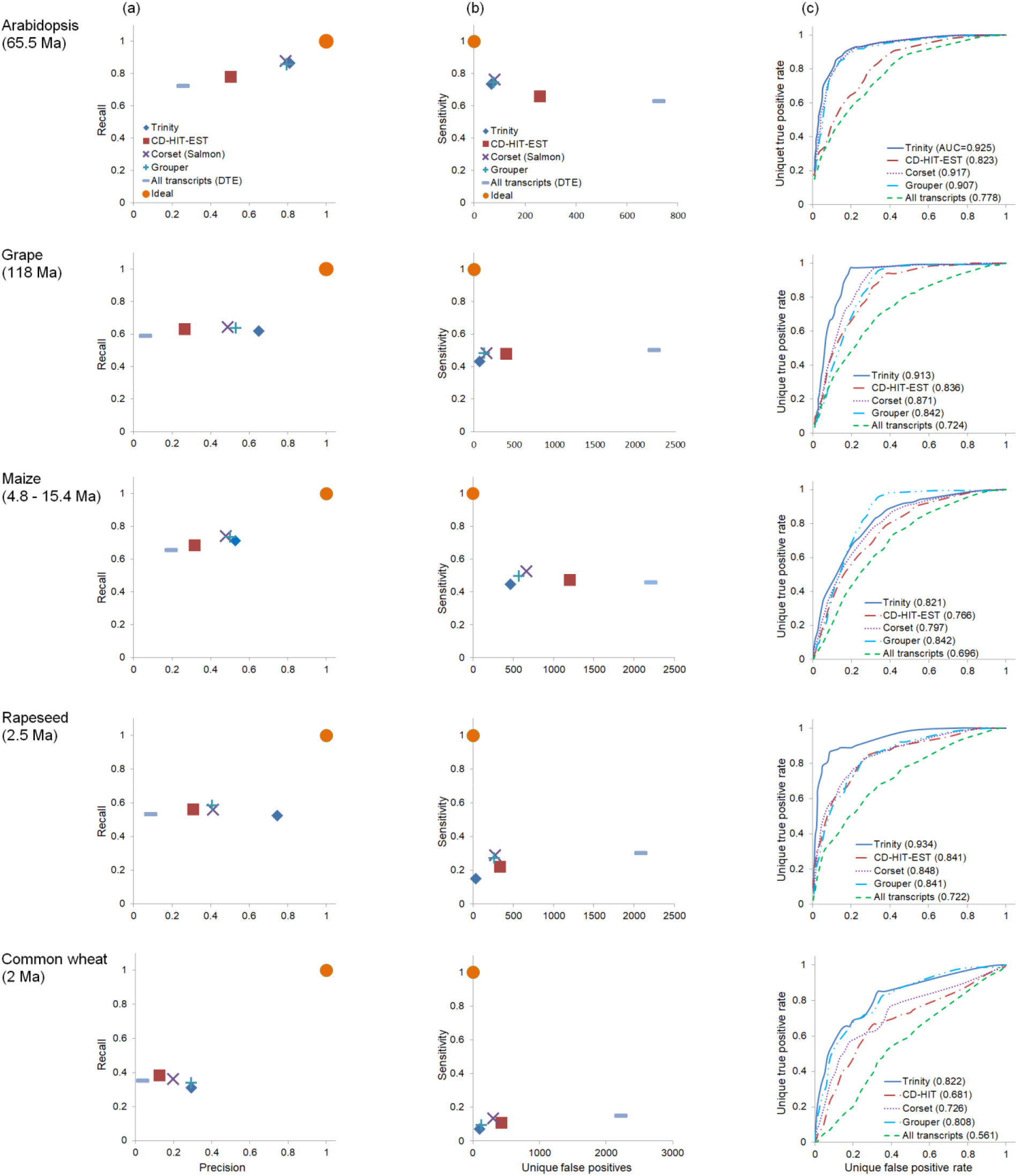
Performance of clustering methods in differential expression analyses. The longest transcript in each cluster was used as the representative and used for read mapping. (a) Recall vs precision. (b) Sensitivity vs number of unique false positives. (c) Receiver operating characteristic (ROC) curve; values in brackets indicates AUC.

Alternatively, representing a gene using all transcripts within a cluster by summarizing reads to gene level generated slightly different results compared to using the longest transcripts as representatives (Supplementary Fig. S3). The two strategies differ in that summation increased the recall of Corset and Grouper in grape but decreasing the recall and precision of Trinity in rapeseed. This can be explained by Corset and Grouper tend to under clustering, which is similar to transcript level analysis. The summation might offset the weakness of under clustering, resulted in increasing of recall. On the other hand, Trinity tends to over clustering. The summation can result in wrong identification when over clustering, which leads to decrease of precision and recall.

Among the five species, *de novo* DGE analyses performed the best in Arabidopsis. Trinity, Corset and Grouper recovered approximately 74% of the refDEGs of Arabidopsis, and FPs accounted for approximately 20% of the deDEGs that could be aligned to reference genes (Supplementary Table S4). Performance of *de novo* DGE analyses in grape is lower than that in Arabidopsis. The poorer performance in grape may be attributed to the higher alternative splicing rate in grape (30%) compared to Arabidopsis (1.2%; Zhang *et al*., 2015) and more frequent recent gene duplications in grape as an evident from more paralogs with *Ks* close to zero compared to Arabidopsis (Supplementary Fig. S2).

DGE analyses performed the worst in rapeseed and wheat, in which Trinity, Corset and Grouper recovered 7–30% of the refDEGs, and FPs accounted for >59% of the deDEGs that could be aligned to reference genes (Supplementary Table S4). Highly similar homeologs produced by the allopolyploidy events led to both low quality of *de novo* assembly (Gutierrez-Gonzalez and Garvin, 2017) and clustering (Davidson and Oshlack, 2014). However, we need to acknowledge that both rapeseed and wheat have extremely complex genomes. Overall, our benchmark study suggested that when the most recent polyploidy event (or split of parental lineages in case of allopolyploidy) happened more than a few million years ago, DGE analysis using *de novo* assembled transcriptomes can be reasonably accurate.

In addition to DGE analysis, clustering is also useful for phylogenomic analysis using *de novo* transcriptome assembly. CD-HIT-EST has been widely used to remove redundancy of *de novo* assembly, e.g. Yang and Smith (2014), which will be further used in orthology inference. According to our analysis, Trinity, Corset and Grouper all perform much better than CD-HIT-EST in the number of correct clusters, precision, and recall. When over clustering and erroneously removing genes is a concern, Corset or Grouper performs the best among the four for downstream phylogenomic analyses, with the proportion of correct clusters being as high as 58.9% in Arabidopsis.

### 3.4 Removing contaminated DEGs is important for downstream functional analysis

Among the five species, Arabidopsis had the highest proportion of TPs among all deDGEs (64.9%), followed by Grape and maize, which had 27.2% and 32% of TPs respectively (Fig. 3). Rapeseed and wheat, however, both had less than 6.6% of TPs. All five species had 1–252 NR-annotated deDEGs, which were not annotated by genome annotation, but annotated by conspecific sequences in NR. This result is consistent with Wang and Gribskov (2017) that *de novo* assembly is beneficial even when a reference genome is available. All five species also have 7.3–89.3% of deDEGs as contamination (Fig. 3), especially rapeseed that was infected by a fungus.

**Figure 3.**
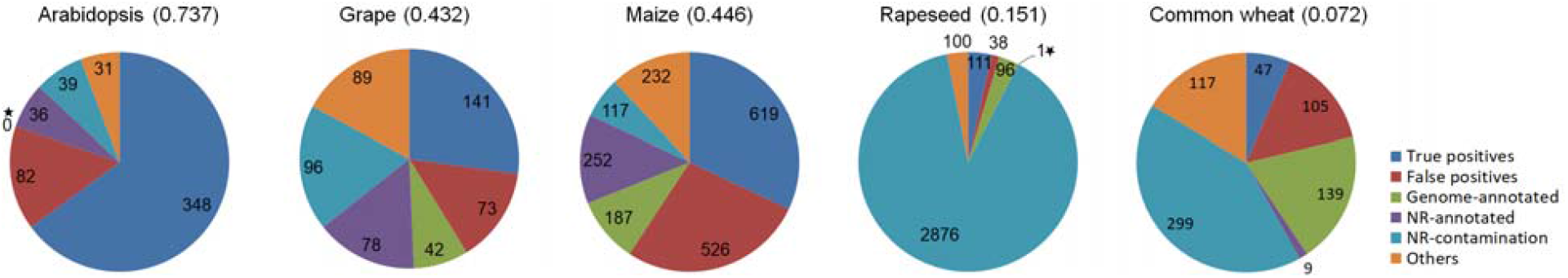
Composition of deDEGs recovered using longest transcripts within Trinity clusters. Sensitivity of DGE analysis was indicated next to species names. Genome-annotated, NR-annotated, and NR-contamination refer to deDEGs that cannot be aligned to any reference gene with our stringent BLAT matching criteria, but can be aligned to reference genome and genes with rnaQUAST, conspecific sequence(s) from the NR database, and sequence(s) of other species in NR respectively. Others includes deDEGs that were mis-assembled (e.g. chimeras), deDEGs that cannot be aligned to any reference gene or NR, and deDEGs that aligned to reference genes, but DESeq2 results lack p-values. The star in the pie of Arabidopsis indicates genome annotated; the star in the pie of rapeseed indicates NR-annotated.

Although contamination may not affect the sensitivity, recall, and precision of DGE analyses (Supplementary results, Supplementary Table S2, and Supplementary Fig. S1), the contaminated DEGs may negatively affect the downstream analyses such as Gene Ontology term enrichment. Moreover, it is more efficient to remove contamination from the relatively small number of DEGs compared to from sequencing reads or all assembled transcripts. Therefore, we recommend verifying the source of DEGs before carrying out further analyses.

### 3.5 Gene co-expression network analyses using *de novo* assembly recovered similar results as reference-based analysis in Arabidopsis

Gene co-expression analysis using *de novo* assembly of Arabidopsis yielded comparable results to the analysis using reference genes. In the remaining four species, however, *de novo* analyses either did not identify any module significantly correlated with the treatment or identified a different module compared to reference-based analysis.

One to two samples for each species were removed given that they were outliers in sample dendrograms (Supplementary Fig. S4). Co-expression network analysis using *de novo* assembled transcriptomes recovered 61 modules in Arabidopsis. Among them, one module (the *“de novo* module”) was significantly correlated with the treatment (correlation-value = 0.920, and p-value = 3.65E-09; Supplementary Table S5). Reference-based analysis recovered 41 modules, among which one module (the “reference module”) was significantly correlated with treatment (correlation-value = 0.903, and p-value = 5.06E-08). The *de novo* and reference modules included 3619 transcripts and 1527 genes respectively. Among the 3619 transcripts, 3475 corresponded to 3269 reference genes. Of the 3475 transcripts, 1395 corresponded to 1249 genes in the reference module, and the remaining 2080 were not in the same reference module (Fig. 4). Among the 2080 transcripts, 4 corresponded to genes in two other reference modules but account for only 1.8% and 3.8% of those two modules respectively.

**Figure 4.**
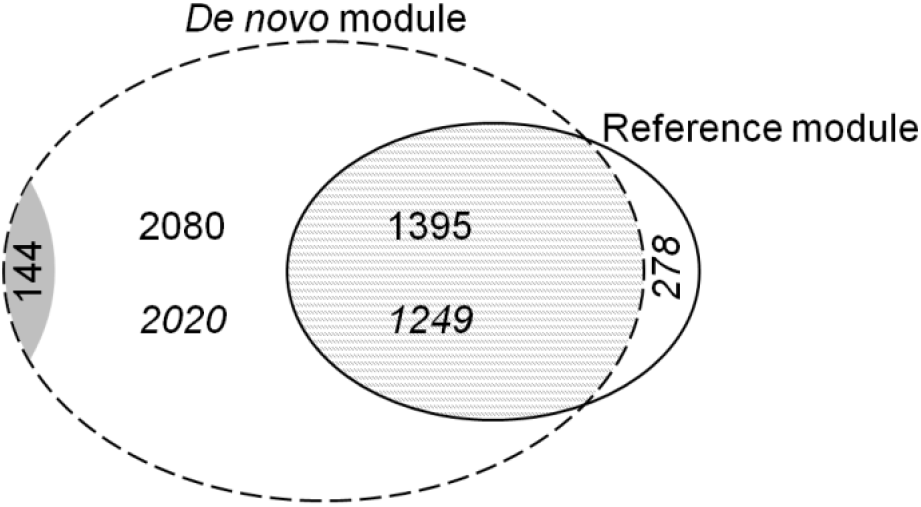
Comparison of the *de novo* module (dashed line) and the reference module (solid line), both of which were significantly correlated with treatment in Arabidopsis. Numbers in normal font indicate the number of transcripts, and numbers in italic indicate corresponding reference genes. The grey area indicates transcripts that do not have any positive match to reference genes.

In maize, 163 *de novo* and 85 reference modules were recovered. One *de novo* module was significantly correlated with the treatment (Supplementary Table S5). The module shared 69 genes with a reference module that included 223 genes. However, this reference module was ranked as the second highest in correlation to treatment. In rapeseed, *de novo* and reference gene-based analysis each yielded one module significantly correlated with treatment separately. However, the *de novo* module and reference module did not form a module pair in either species. In grape and wheat, *de novo* based analysis yielded one module significantly correlated with treatment or traits separately. However, no module was significantly correlated with treatment or traits in reference gene-based analysis.

In addition to quality of transcript assembling and putative gene clustering, experimental design and the number of datasets are important. None of the datasets we used were initially designed for co-expression network analysis except the wheat dataset. Among the five species, Arabidopsis is the only one with the recovery stage sampled, in addition to the control and three additional time points during stress treatment. In co-expression network analysis, a higher number of samples and data points usually lead to more robust and refined results, and at least 20 samples were recommended by WGCNA (Langfelder and Horvath, 2008). However, publicly available RNA-seq datasets with sufficient time points and biological replicates are scarce, and we were only able to include 12 samples for both grape and maize. We would like to emphasize that with proper experimental design (stress with recovery with a sufficient number of time points), and genome with low complexity (e.g. polyploidization not too recent), *de novo* assembled transcriptome can potentially recover the correct module.

## 4 Conclusion

Our analyses show that unless the polyploidy event is as recent as a few million years ago or less, both DGE and co-expression network analyses can be a powerful tool in diverse organisms without reference genomes. By benchmarking performance of four clustering approaches in five species of different polyploidy history, we demonstrated that CD-HIT-EST is actually not the best for clustering while it is heavily used. We would suggest using Trinity clusters and picking longest transcripts for DGE analysis if a study aims to obtain fewer but more reliable DEGs; using Corset or Grouper clusters and summarizing reads to gene level for DGE analysis if a study aims to obtain a broader list of DEGs. Furthermore, we demonstrated the advantage of DGE analysis using assembled transcripts in that it is capable of recovering differentially expressed genes that are missing from the reference genome assembly or annotation. On the other hand, the prevalence of contamination in DGE results points to the importance of verifying the source of transcripts before carrying out functional annotation and interpreting the results. Finally, our analyses highlight the importance of experimental design and a sufficient number of data points and biological replicates in co-expression network analysis, especially when a reference genome is not available.

With the improved sequencing power (e.g. Illumina NovaSeq) and long-read transcriptome sequencing (e.g. IsoSeq from PacBio, and Nanopore technologies), gene expression analysis, both DGE and co-expression network are expected to rapidly expand functional genomic research into diverse organisms with new analytical powers. Our benchmark analyses provided the bases for continued phylotranscriptomics, DGE, and gene co-expression network analyses in diverse organisms without reference genomes. When combined with a phylogenetic sampling of gene expression experiments, the power can be further improved (Dunn *et al*., 2018), leading to the discovery of genes and modules underline evolutionary novelties in diverse organisms.

## Acknowledgements

We thank Minnesota Supercomputing Institute (MSI) for providing computing resource; Guan-Jing Hu for discussion on gene co-expression network analysis; Nadia M Davidson, Avi Srivastava, and Laraib Malik for assistance on data analyses; Michael I. Love for assistance on DESeq2; Peter Langfelder and Wen Zhou for assistance on WGCNA; Andrey Prjibelski for assistance on rnaQUAST; Joël Lafond Lapalme for assistance on MCSC and Lian-Fu Chen for assistance on software installation.

## References

Arias, T. et al. (2014) Diversification times among Brassica (Brassicaceae) crops suggest hybrid formation after 20 million years of divergence. Am. J. Bot., 101, 86–91.

Beilstein, M.A. et al. (2010) Dated molecular phylogenies indicate a Miocene origin for *Arabidopsis thaliana*. Proc. Natl. Acad. Sci. USA., 107, 18724–18728.

Berthelot, C. et al. (2014) The rainbow trout genome provides novel insights into evolution after whole-genome duplication in vertebrates. Nat. Commun., 5, 3657.

Blanc, G. and Wolfe, K.H. (2004) Widespread paleopolyploidy in model plant species inferred from age distributions of duplicate genes. Plant Cell, 16, 1667–1678.

Bolger, A.M., Lohse, M. and Usadel, B. (2014) Trimmomatic: a flexible trimmer for Illumina sequence data. Bioinformatics, 30, 2114–2120.

Bushmanova, E. et al. (2016) rnaQUAST: a quality assessment tool for *de novo* transcriptome assemblies. Bioinformatics, 32, 2210–2212.

Camacho, C. et al. (2009) BLAST+: architecture and applications. BMC Bioinformatics, 10, 421.

Carruthers, M. et al. *De novo* transcriptome assembly, annotation and comparison of four ecological and evolutionary model salmonid fish species. BMC Genomics 2018, 19, 32.

Coolen, S. et al. (2016) Transcriptome dynamics of Arabidopsis during sequential biotic and abiotic stresses. Plant J., 86, 249–267.

Davidson, N.M. and Oshlack, A. (2014) Corset: enabling differential gene expression analysis for *de novo* assembled transcriptomes. Genome Biol., 15, 410.

Dobin, A. et al. (2013) STAR: ultrafast universal RNA-seq aligner. *Bioinformatics*, 29, 15–21. Fawcett, T. (2006) An introduction to ROC analysis. Pattern Recog. Lett., 27, 861–874.

Feldmann, K.A. and Goff, S.A. (2014) The first plant genome sequence—*Arabidopsis thaliana*. Adv. Bot. Res., 69, 91–117.

Forest, F. and Chase, M.W. (2009) Eudicots. In: Hedges, S.B. and Kumar, S., editors, The Timetree of Life, pp. 169–176. Oxford University Press, New York, NY.

Garcia, K. et al. (2017) Physiological responses and gene co-expression network of mycorrhizal roots under K+ deprivation. Plant Physiol., 173, 1811–1823.

Grabherr, M.G. et al. (2011) Full-length transcriptome assembly from RNA-Seq data without a reference genome. Nat. Biotechnol., 29, 644–652.

Gutierrez-Gonzalez, J.J. and Garvin, D.F. (2017) De novo Transcriptome assembly in polyploid species. In: Gasparis, S., editor, Oat Methods and Protocols, pp. 209–221. Humana Press, New York, NY.

Heyduk, K. et al. (2018) Shifts in gene expression profiles are associated with weak and strong Crassulacean acid metabolism. Am. J. Bot., 105, 587–601.

Honaas, L.A. et al. (2016) Selecting superior *de novo* transcriptome assemblies: lessons learned by leveraging the best plant genome. PLoS One, 11, e0146062.

Jiang, J. et al. (2017) Integrating omics and alternative splicing reveals insights into grape response to high temperature. Plant Physiol., 173, 1502–1518.

Kent, W.J. (2002) BLAT—the BLAST-like alignment tool. Genome Res., 12, 656–664.

Lafond-Lapalme, J. et al. (2017) A new method for decontamination of *de novo* transcriptomes using a hierarchical clustering algorithm. Bioinformatics, 33, 1293–1300.

Langfelder, P. and Horvath, S. (2008) WGCNA: an R package for weighted correlation network analysis. BMC Bioinformatics, 9, 559.

Langmead, B. and Salzberg, S.L. (2012) Fast gapped-read alignment with Bowtie 2. Nat. Methods, 9, 357.

Lee, T.H. et al. (2012) PGDD: a database of gene and genome duplication in plants. Nucleic Acids Res., 41, D1152–D1158.

Lee, T.H. et al. (2017) Plant genome duplication database. Methods Mol, Biol., 1533, 267–277.

Li, B. and Dewey, C.N. (2011) RSEM: accurate transcript quantification from RNA-Seq data with or without a reference genome. BMC Bioinformatics, 12, 323.

Li, W. and Godzik, A. (2006) Cd-hit: a fast program for clustering and comparing large sets of protein or nucleotide sequences. Bioinformatics, 22, 1658–1659.

Li, Z. et al. (2018) Multiple large-scale gene and genome duplications during the evolution of hexapods. Proc. Natl. Acad. Sci. USA., 115, 4713–4718.

Loraine, A. (2009) Co-expression analysis of metabolic pathways in plants. Methods Mol. Biol., 553, 247–264.

Love, M.I., Huber, W. and Anders, S. (2014) Moderated estimation of fold change and dispersion for RNA-seq data with DESeq2. Genome Biol., 15, 550.

Ma, J. et al. (2017) Transcriptomics analyses reveal wheat responses to drought stress during reproductive stages under field conditions. Front. Plant Sci., 8, 592.

Malik, L., Almodaresi, F. and Patro, R. (2018) Grouper: Graph-based clustering and annotation for improved *de novo* transcriptome analysis. Bioinformatics, 1, 8.

Marcet-Houben, M. and Gabaldón, T. (2015) Beyond the whole-genome duplication: phylogenetic evidence for an ancient interspecies hybridization in the baker’s yeast lineage. PLoS Biol., 13, e1002220.

Middleton, C.P. et al. (2014) Sequencing of chloroplast genomes from wheat, barley, rye and their relatives provides a detailed insight into the evolution of the Triticeae tribe. PLoS One, 3, e85761.

Mimura, M. et al. (2016) Arabidopsis TH2 encodes the orphan enzyme thiamin monophosphate phosphatase. Plant Cell, 28, 2683–2696.

Misof, B. et al. (2014) Phylogenomics resolves the timing and pattern of insect evolution. Science, 346, 763–767.

Nakasugi, K. et al. (2014) Combining transcriptome assemblies from multiple *de novo* assemblers in the allo-tetraploid plant *Nicotiana benthamiana*. PLoS One, 9, e91776.

Patro, R. et al. (2017) Salmon provides fast and bias-aware quantification of transcript expression. Nat. Methods, 14, 417–419.

Roberts, W.R. and Roalson, E.H. (2017) Comparative transcriptome analyses of flower development in four species of *Achimenes* (Gesneriaceae). BMC Genomics, 18, 240.

Schulz, M.H. et al. (2012) Oases: robust *de novo* RNA-seq assembly across the dynamic range of expression levels. Bioinformatics, 28, 1086–1092.

Soneson, C., Love, M. and Robinson, M. (2015) Differential analyses for RNA-seq: transcript-level estimates improve gene-level inferences. F1000Research, 4, 1521.

Song, L. and Florea, L. (2015) Rcorrector: efficient and accurate error correction for Illumina RNA-seq reads. Gigascience, 4, 48.

Srivastava, A. et al. (2016) Accurate, fast and lightweight clustering of *de novo* transcriptomes using fragment equivalence classes. BioRxiv, 1604.03250.

Swigoftová, Z. et al. (2004) Close split of sorghum and maize genome progenitors. Genome Res., 14, 1916–1923.

Teng, M. et al. (2016) A benchmark for RNA-seq quantification pipelines. Genome Biol., 17, 74.

van Rijsbergen, C.J. (1979) Information Retrieval, University of Glasgow, PhD thesis, London.

Wang, S. and Gribskov, M. (2017) Comprehensive evaluation of *de novo* transcriptome assembly programs and their effects on differential gene expression analysis. Bioinformatics, 33, 327–333.

Wang, Y. et al. (2017) Transcriptome association identifies regulators of wheat spike architecture. Plant Physiol., 175, 746–757.

Wu, J. et al. (2016) Comparative transcriptomic analysis uncovers the complex genetic network for resistance to *Sclerotinia sclerotiorum* in *Brassica napus*. Sci. Reports, 6, 19007.

Yang, Y. and Smith, S.A. (2013) Optimizing *de novo* assembly of short-read RNA-seq data for phylogenomics. BMC Genomics, 14, 328.

Yang, Y. and Smith, S.A. (2014) Orthology inference in nonmodel organisms using transcriptomes and low-coverage genomes: improving accuracy and matrix occupancy for phylogenomics. Mol. Biol. Evol., 31, 3081–3092.

Yang, Y. et al. (2015) Dissecting molecular evolution in the highly diverse plant clade Caryophyllales using transcriptome sequencing. Mol. Biol. Evol., 32, 2001–2004.

Zhang, C., Yang, H. and Yang, H. (2015) Evolutionary character of alternative splicing in plants. Bioinform. Biol. Insights, 9(Suppl 1), 47–52.

